# Social experience alters oxytocinergic modulation in the nucleus accumbens of female prairie voles

**DOI:** 10.1101/2021.07.06.451323

**Authors:** Amélie M. Borie, Sena Agezo, Parker Lunsford, Arjen J. Boender, Ji-Dong Guo, Hong Zhu, Gordon J. Berman, Larry J. Young, Robert C. Liu

## Abstract

Social relationships are dynamic and evolve with shared and personal experiences. Whether the functional role of social neuromodulators also evolves with experience to shape the trajectory of relationships is unknown. We utilized pair bonding in the socially monogamous prairie voles as an example of socio-sexual experience that dramatically alters behaviors displayed toward other individuals. We investigated oxytocin-dependent modulation of excitatory synaptic transmission in the nucleus accumbens as a function of pair bonding status. We found that an oxytocin receptor agonist decreases the amplitude of spontaneous Excitatory Postsynaptic Currents (EPSCs) in sexually naive virgin, but not pair-bonded, female voles, while it increases the amplitude of electrically evoked EPSCs in paired voles, but not in virgins. This oxytocin-dependent potentiation of synaptic transmission relies on the *de novo* coupling between oxytocin receptor signaling and endocannabinoid CB1 receptor signaling in pair bonded voles. Blocking CB1 receptors after pair bond formation increases the occurrence of a specific form of social rejection – defensive upright response – that is displayed towards the partner but not towards a novel individual. Altogether, our results demonstrate that oxytocin’s action in the nucleus accumbens is changed through social experience in a way that regulates the trajectory of social interactions as the relationship with the partner unfolds, potentially promoting the maintenance of a pair bond by inhibiting aggressive responses. These results provide a mechanism by which social experience and context shift oxytocinergic signaling to impact neural and behavioral responses to social cues.

## Introduction

Social relationships evolve over time as interactions – both positive and negative – reshape how one subsequently perceives (Suvilehto et al. 2015), engages with (Walum and Young 2018) and feels (Whitehouse et al. 2014) about another individual. Thus, the brain mechanisms acting during the formation of a relationship and its later expression and maintenance are likely to change over the course of subsequent encounters between two individuals. While considerable work has helped elucidate the neural mechanisms underlying the formation and expression of social relationships (Walum and Young 2018; Amadei et al. 2017; Smith, Lei, and Wang 2013), much less is known about what modulates their maintenance. In particular, although neuromodulators are well documented to act centrally in the brain to modulate social behaviors (Miranda and Liu 2009; Banerjee and Liu 2013; Donaldson and Young 2008), here we investigated whether that action can change over the course of a social relationship.

Oxytocin (OXT) is a key neuromodulator for the regulation of social behaviors (Froemke and Young 2021; Johnson and Young 2017), and a target for the treatment of neuropsychiatric disorders (Meyer-Lindenberg et al. 2011; DeMayo et al. 2019; Ford and Young 2021). OXT is produced mainly by neurons located in the supra-optic and paraventricular nuclei of the hypothalamus. OXT is released within the brain as relationships form and facilitate experience-dependent social adaptations including the formation of social memories (Ferguson et al. 2001; Lukas et al. 2013), the establishment of mating-induced pair bonds (Walum and Young 2018) and maternal responsiveness (Marlin et al. 2015). It is believed that OXT action results in modulation of neuronal activity in brain areas regulating social behaviors (Johnson et al. 2017) to increase the salience and reinforcing value of social stimuli (Young 2015; Shamay-Tsoory and Abu-Akel 2016). Furthermore, OXT neurons are activated in response to social stimuli including social touch (Tang et al. 2020), infant cries (Valtcheva et al. 2021) and mating. These events, which occur as the relationship develops, offer an opportunity for the OXT system to act after the initial encounter to modulate the maintenance of the social bonds.

Elucidating whether OXT’s actions may change over the course of a social bond requires a model where animals form long term relationships. Prairie voles (*Microtus ochrogaster*) are rodents that, upon social interaction and mating, form a lifelong pair bond resulting in a dramatic change in the way they interact with other individuals (McGraw and Young 2010). These behavioral shifts are accompanied by concomitant shifts in neural dynamics within brain networks important for the regulation of social behavior (Scribner et al. 2020; Amadei et al. 2017), particularly involving a brain area at the intersection between the social neural network and the mesolimbic reward system: the nucleus accumbens (NAc). Interestingly, prairie voles and other monogamous species, including marmosets and human express high levels of oxytocin receptors in the NAc while non-monogamous species, such as laboratory mice and rhesus macaques, do not (Young and Zhang 2021; Freeman and Young 2016; Bethlehem et al. 2017). Furthermore, OXTR signaling in the NAc plays a critical role for the formation of a preference for a known individual, and the density of NAc OXTR correlates with the preference (Young et al. 2001; Keebaugh et al. 2015; Keebaugh and Young 2011). Thus, OXTR signaling in the NAc may provide mechanistic insights into experience and context dependent modulation of social behavior.

Here, we studied OXT-dependent modulation of NAc core activity both in sexually naïve animals and in pair bonded females to determine whether OXT may have different physiological effects in the context of pair bond formation versus maintenance. Using a combination of slice physiology, pharmacology, CRISPR genome editing and behavior, we provide evidence that OXT may reduce synaptic noise in the NAc of virgin animals, potentially facilitating bonding, yet potentiates glutamatergic transmission in pair bonded females by coupling to an endocannabinoid receptor (CB1) dependent, presynaptic mechanism. Furthermore, by blocking CB1 *in vivo*, we demonstrate that this mechanism specifically represses defensive responses in the partner’s presence, which presumably helps maintain a trajectory to consolidate the pair bond. Altogether, we demonstrate that the action of neuromodulators such as OXT are context dependent and should be considered in light of the specific social history of an individual.

## Results

### The effect of OXT on spontaneous and evoked synaptic activity depends on social experience

We sought to examine the electrophysiological effects of OXT on NAc core medium spiny neurons (MSNs) and whether they are influenced by social experience. We ovariectomized sexually naive female subjects and cohabitated them with either a same sex littermate (Virgin group) or a male (Paired group) for 24h. We performed slice electrophysiology recordings of MSNs in whole cell configuration 48h later (Figure 1A). Basal properties of NAc MSNs remained the same in both Virgin and Paired groups (Supplemental Figure 1) and were consistent with a prior study in voles (Willett et al. 2018). We then applied in the bath a high specificity OXTR agonist TGOT (10^-7^M) (Chini, Manning, and Guillon 2008) for 10-min. TGOT did not influence the frequency of spontaneous Excitatory PostSynaptic Currents (sEPSC) in either Virgin or Paired females (Figure 1B). However, TGOT induced a significant decrease of the sEPSCs amplitude in Virgin but not in Paired females (Figure 1C-D), potentially decreasing the effect of spontaneous excitation in Virgins.

**Figure 1:**
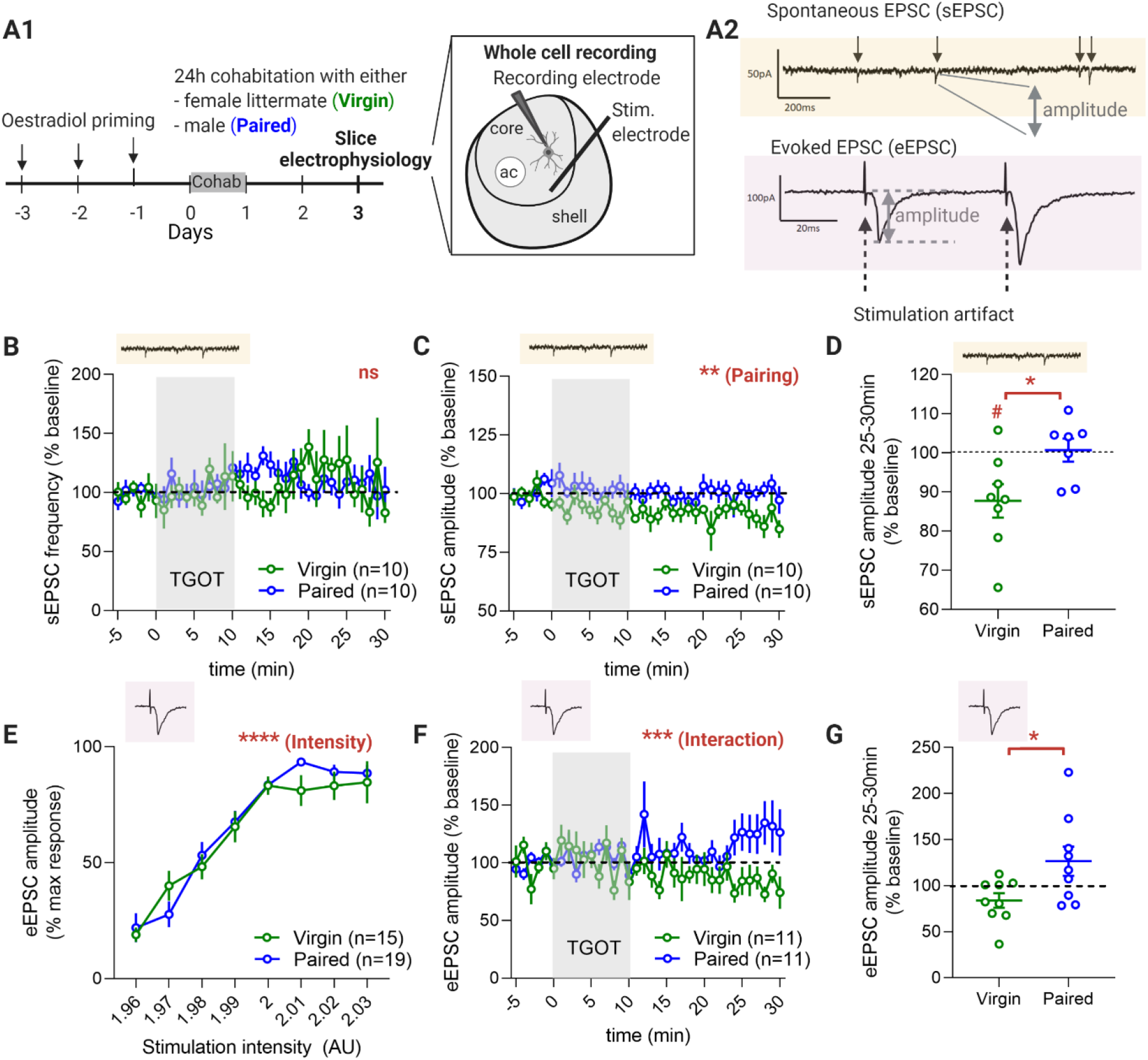
Pair bonding differentially alters spontaneous and evoked synaptic activity in response to TGOT. **(A1)** Experimental timeline (left) and schematic showing the position of the stimulation and recording electrodes (right). **(A2)** Representative examples of spontaneous EPSCs recordings (top, each arrow points to a synaptic event) and of evoked EPSC (bottom, dotted arrow point toward the stimulation artefacts and grey double arrow indicates the measured amplitude). **(B)** Effect of 10min bath application of TGOT on spontaneous EPSC frequency. Data (mean ± SEM) expressed as % relative to baseline. Two-way analysis of variance (ANOVA): effect of time F(35, 522)=1.042, p=0.4054; effect of Social experience (Social exp.). F(1,18)=0.2313, p=0.636; interaction F(35,522)=1.078, p=0.352. **(C)** Effect of 10min bath application of TGOT on spontaneous EPSC amplitude. Data (mean ± SEM) expressed as % relative to baseline. Two-way ANOVA: effect of time F(6.206, 92.03)=0.8964, p=0.504; effect of Social exp. F(1,18)=8.552, p=0.009; interaction F(35,519)=0.9250, p=0.595. **(D)** Average over a 5min period of the effect of 10min bath application of TGOT on spontaneous EPSC amplitude. Data (mean ± SEM) expressed as % relative to the baseline. Arbitrary Unit (AU). Unpaired t-test, p=0.0306. One sample Wilcoxon test compared to 100%: Virgin p=0.023; Paired p=0.812. **(E)** Amplitude of the evoked EPSC as a function of stimulation intensity. Data (mean ± SEM) expressed as % relative to the maximum amplitude. Two-way ANOVA: effect of stimulation intensity F(7, 204)=66.39, p<0.0001; effect of Social exp. F(1,32)=0.4529, p=0.506; interaction F(7,204)=1.232, p=0.286. **(F)** Time-course of the effect of 10min bath application of TGOT on evoked EPSC amplitude. Data (mean ± SEM) expressed as % relative to the baseline. Two-way ANOVA: effect of time F(6.674,124.7)=1.301, p=0.257; effect of Social exp. F(1,20)=3.631, p=0.071; interaction F(35,654)=2.247, p<0.0001. **(G)** Average over a 5min period of the effect of 10min bath application of TGOT on evoked EPSC amplitude. Data (mean ± SEM) expressed as % relative to the baseline. Unpaired t-test, p=0.028. Number of cells as indicated in each panel.

To further examine TGOT’s effects on NAc neurotransmission, we electrically evoked EPSCs (eEPSC) in whole cell recordings. The relationship between the stimulation intensity and the eEPSC amplitude was not influenced by the social experience of the animals (Figure 1E), indicating a similar sensitivity of neurons across groups to the electrical stimulation. Moreover, the raw amplitudes of the eEPSCs were not different between groups, suggesting that social experience did not generally increase the basal excitation of MSNs (Supplemental Figure 1H). Strikingly, there was a significant interaction between the social experience of the animal and time (Figure 1F). TGOT produced a more potentiated response in Paired compared to Virgin females (Figure 1G), potentially increasing excitatory neurotransmission in Paired females.

### The amplitude of OXT-induced potentiation correlates with the strength of the social preference in field recordings

Because the effect of TGOT on evoked glutamatergic activity depends on social experience, we next investigated whether the strength of the post-cohabitation partner preference was correlated to the amplitude of this electrophysiological response. We performed modified partner preference tests – social preference tests (SPT) – before the slice electrophysiology experiment to evaluate the pair bond strength (Figure 2A-B). Males were placed under cups, restricting physical contact with the experimental female. We assessed the time spent near either male. Before the 24h cohabitation, there was no consistent preference in either the Virgin or Paired group. After cohabitation, Virgin females still showed no systematic preference, but those cohabitated with a male spent significantly more time near that individual (Figure 2C). Hence, like the traditional partner preference test, the SPT reflects the strength of the animal’s pair bond.

**Figure 2:**
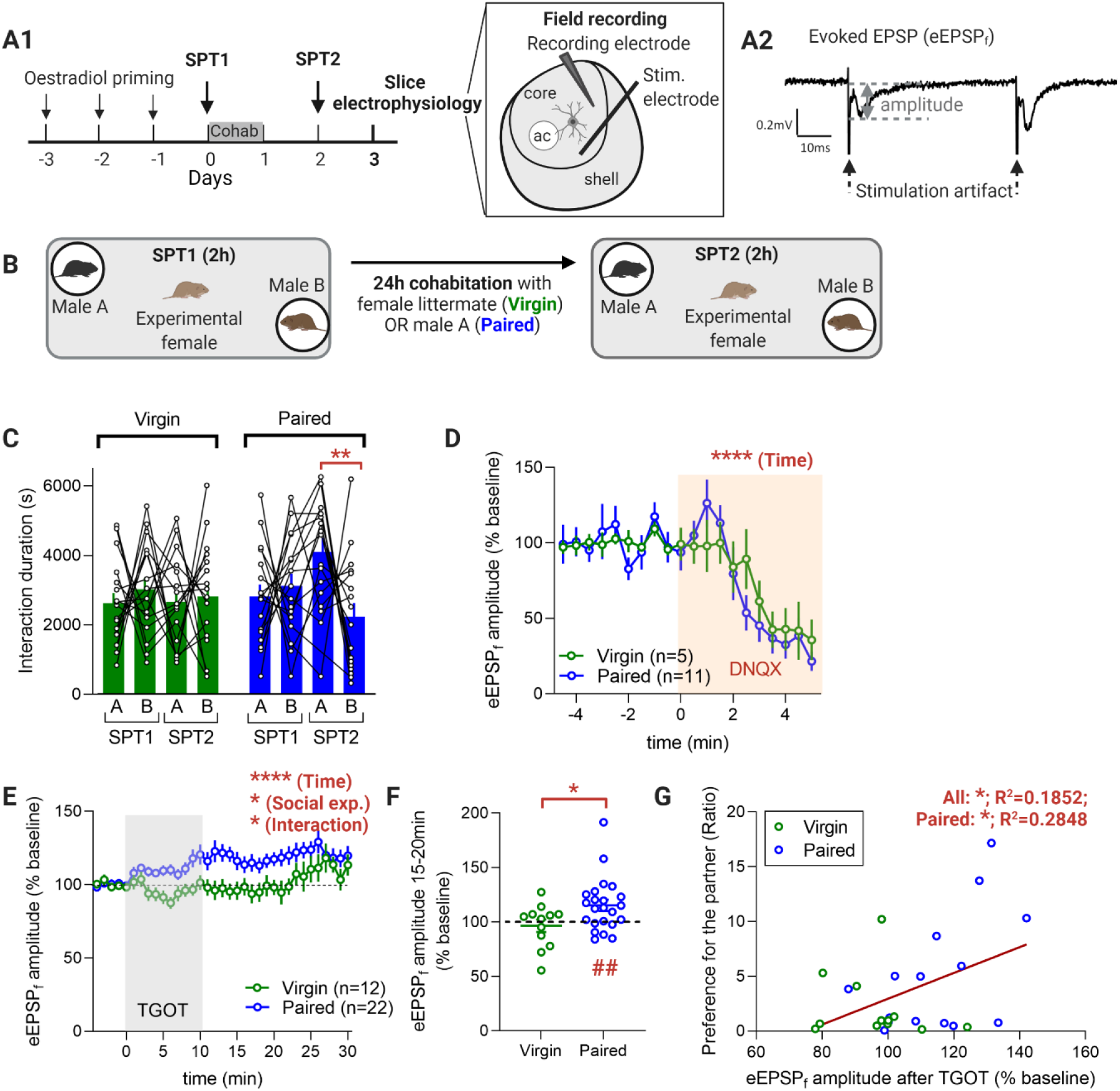
The amplitude of oxytocin-induced potentiation in field recordings correlates with the strength of the social preference. **(A1)** Experimental timeline (left) and schematic showing the position of the stimulation and recording electrodes (right). **(A2)** representative example of evoked EPSP (right, dotted arrow point toward the stimulation artefact and grey double arrow indicates the measured amplitude). **(B)** Experimental protocol – Social preference test (SPF). **(C)** Duration of the interactions with male A and male B in SPF1 and SPT2 for Virgin and Paired individuals. Data (mean ± SEM), n=13 voles for SFP1 Virgin, 12 for SPF2 Virgin, 14 for SPF1 Paired and 14 for SPF2 Paired. Three-way ANOVA: effect of Social exp. F(1,126)=1.273, p=0.261; SPT1 vs SPT2 F(1,126)=0.04291, p=0.836; A vs B F(1,126)=0.9981, p=0.320; interaction Social exp. × SPT1 vs SPT2 F(1,126)=0.3026, p=0.583; interaction Social exp. × A vs B F(1,126)=4.403, p=0.038; interaction SPT1 vs SPT2 × A vs B F(1,126)=5.611, p=0.019; interaction Social exp. × SPT1 vs SPT2 × A vs B F(1,126)=3.619, p=0.059; Post-hoc Sidak test (A vs B): SPT1 Virgin p=0.999, SPT2 Virgin p>0.999, SPT1 Paired p>0.999, SPT2 Paired p=0.004. **(D)** Effect of DNQX on the evoked EPSP amplitude (field recording). Data (mean ± SEM) expressed as % relative to baseline, n as indicated. Two-way ANOVA: effect of time F(19,266)=8.977, p<0.0001; effect of Social exp. F(1,14)=0.09597, p=0.761; interaction F(19,266)=0.5327, p=0.947. **(E)** Time course of the effect of 10min bath application of TGOT on evoked EPSP amplitude (field recording). Data (mean ± SEM) expressed as % relative to baseline, n as indicated. Two-way ANOVA: effect of time F(34,1083)=3.399, p<0.0001; effect of Social exp. F(1,32)=6.888, p=0.013; interaction F(34,1083)=1.536, p=0.026. **(F)** Average over a 5min period of the effect of 10min bath application of TGOT on evoked EPSP amplitude (field recording). Data (mean ± SEM) expressed as % relative to baseline. Unpaired t-test, p=0.0290. One sample Wilcoxon test compared to 100%: Virgin p=0.9097, Paired p=0.004. **(G)** Correlation between the amplitude of the effect of TGOT on the evoked EPSP amplitude during the 2 last minutes of TGOT application and the preference for the partner. Pearson correlation coefficient as indicated and R^2^=0.03412 for Virgin. All: p=0.025, Paired: p=0.040, Virgin: p=0.566. Linear regression (All): y=0.1177×74.92.

To relate this behavioral preference to the electrophysiological response to TGOT, we turned to field recordings. As a reflection of activity in a larger population of neurons, the evoked Excitatory Postsynaptic Potential in the field (eEPSP_f_) is less sensitive to cell-to-cell variability inherent in whole-cell recordings. The eEPSPf was blocked by DNQX, thus demonstrating its glutamatergic nature. Consistent with the results obtained in whole cell recording, TGOT application differentially modulated the eEPSP_f_ in Virgin and Paired females, as reflected in a significant interaction between group and time (Figure 2E-F). Specifically, the amplitude of the eEPSP_f_ was significantly increased following TGOT application in Paired females, whereas in Virgin females it remained unchanged (Figure 2F). Furthermore, the amplitude of the response correlated with the preference for the partner, defined as the time spent near the partner relative to the stranger during the second SPT (Figure 2G). Therefore, how the NAc responds to OXT depends on the effect of the vole’s prior social experience, pointing to a behavioral relevance of the plasticity in the OXT system’s physiological effects. We then focused on this robust experience-dependent potentiation in Paired females to decipher the underlying mechanism.

### OXT-dependent potentiation in Paired female voles relies on postsynaptic OXTR and a presynaptic mechanism

Since the OXT-dependent potentiation in whole cell recordings was observed in Paired animals for the evoked, but not the spontaneous EPSC amplitude, even when both responses were measured from the same MSN, we suspected the involvement of a pre-synaptic, rather than a post-synaptic, mechanism. To further evaluate this possibility, we measured the effect of TGOT on the Paired pulse ratio (PPR) in evoked EPSC and EPSPf, from whole cell (Figure 3B-C) and field recording (Figure 3D-E) configurations, respectively. The PPR is sensitive to the probability of presynaptic vesicle release (Dobrunz and Stevens 1997). In Paired animals, the stronger the evoked, TGOT-modulated postsynaptic response to the first stimulation pulse in a pair, the smaller the second response and PPR, indicating that TGOT facilitated a larger initial presynaptic release of glutamate, In Virgin animals where there was no systematic TGOT-mediated potentiation, these values were not correlated. Hence, pair bonding experience was sufficient for TGOT to systematically affect presynaptic glutamate release in the NAc.

**Figure 3:**
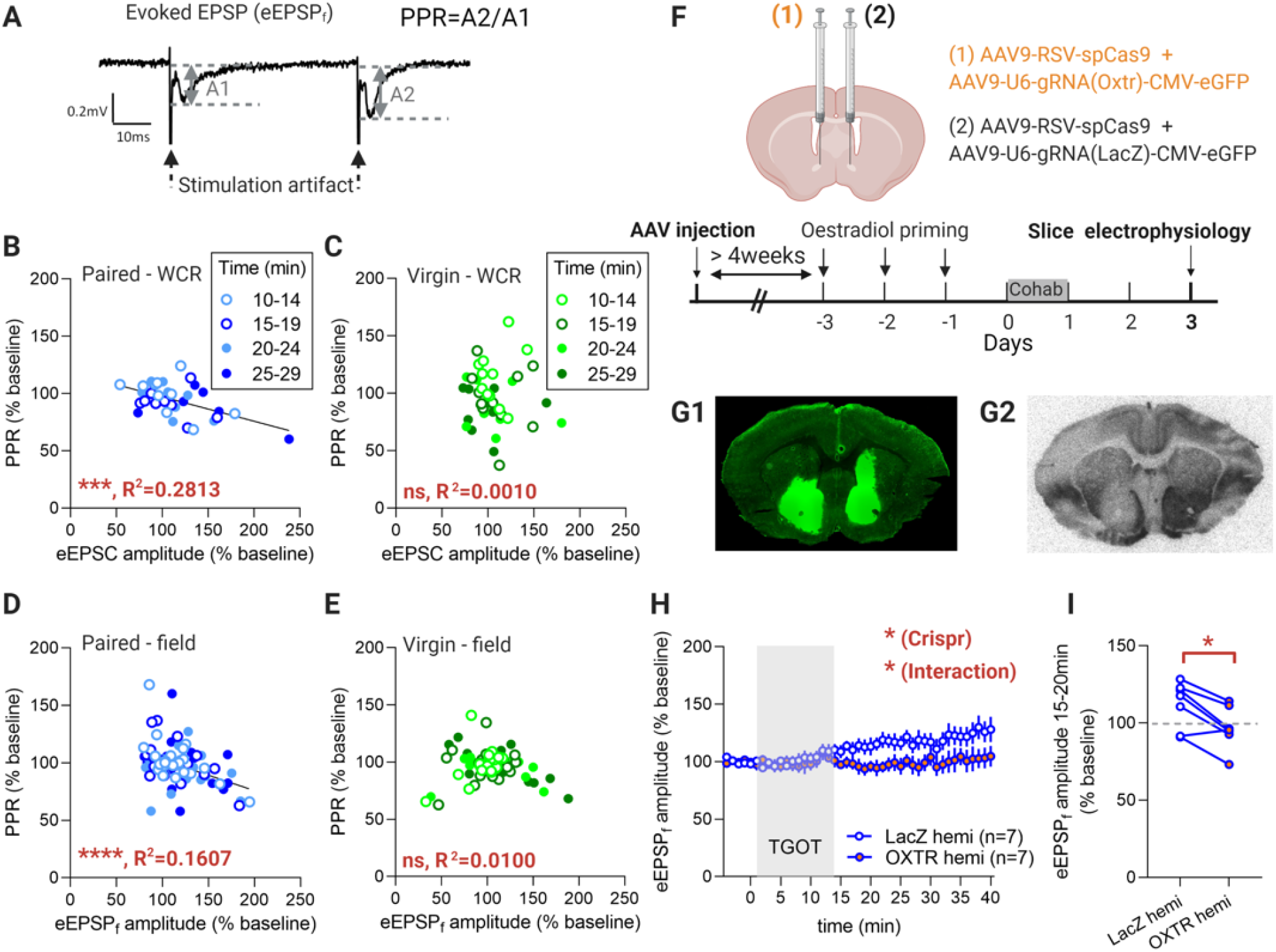
Oxytocin-dependent potentiation in Paired prairie voles relies on a presynaptic mechanism but involves OXTR in NAc cells. **(A)** Representative example of evoked EPSPs (right, dotted arrow point toward the stimulation artefact and grey double arrow indicates the measured amplitude) and formula for Paired pulse ratio (PPR) calculation. **(B)** Correlation between EPSC amplitude and Paired pulse ratio (PPR) in whole cell recordings (WCR) from pair bonded voles. Pearson correlation coefficient as indicated, p=0.0005. Linear regression: y=−0.2128x+556.7. **(C)** Correlation between EPSC amplitude and PPR in WCR from Virgin animals. Pearson correlation coefficient as indicated, p=0.845. **(D)** Correlation between EPSP amplitude PPR in field recording from pair bonded animals. Pearson correlation coefficient as indicated, p<0.0001. Linear regression: y=−0.2676x+483.7. **(E)** Correlation between EPSP amplitude and PPR in field recordings from Virgin animals. Pearson correlation coefficient as indicated, p=0.417. **(B) to (E)** Each dot represents the average over a 5min period (10-14min, 15-19min, 20-24min or 25-29min – timing as indicated in figure 1E or 2E) and for one cell. **(F)** Viral-vector mediated Crispr strategy (top) and timeline of the experiment (bottom). **(G)** Representative examples of the infection visible through the green fluorescence (left) and corresponding autoradiograms (right) showing that the AAV-mediated CRISPR strategy is efficient to reduce OXTR binding in the NAc specifically on the side infected with the sg-RNA targeting OXTR. **(H)** Timeline of the effect of a 10min TGOT application on the evoked EPSP amplitude in NAc knocked down for OXTR and in the contralateral control side (field recordings). Data (mean ± SEM) expressed as % relative to baseline, n as indicated. Two-way ANOVA (data from ipsilateral and contralateral side are Paired) : effect of time F(2.572,15.43)=1.671, p=0.217; effect of the AAV-mediated Crispr strategy F(1.000,6.000)=6.951, p=0.039; interaction F(3.230,19.38)=3.893, p=0.023. **(I)** Average over a 5min period of the effect of a 10min TGOT application on the evoked EPSP amplitude in NAc knocked down for OXTR and in the contralateral control side (field recordings). Data expressed as % relative to baseline. Wilcoxon test, p=0.031.

These results might suggest the involvement of presynaptic OXTR rather than OXTRs located on cells in the NAc. To explicitly test the necessity of NAc OXTR for the OXT-evoked potentiation observed in Paired animals, we used an AAV-mediated CRISPR/Cas9 strategy to knock down OXTR expression in cells from the NAc. If OXTR required for the potentiation are located on the pre-synapse, knocking down OXTR expression in NAc cells should not affect the amplitude of the potentiation. Females were administered a combination of AAV-vectors driving expression of spCas-9 and an sgRNA targeting OXTR in one hemisphere’s NAc and vectors enabling for expression of spCas-9 and a control sgRNA targeting LacZ in the contralateral NAc (Figure 3F), as previously described (Boender and Young 2020). After allowing at least 4 weeks for expression and genome editing, animals were cohabitated with a male for 24h, and their brains were extracted and sliced to perform slice electrophysiology in field recording configuration.

Expression of the AAV coding for the sg-RNA was evaluated (Green fluorescence) and found comparable on both sides (Figure 3G1). Furthermore, autoradiograms performed using I^125^-OVTA show that this strategy was efficient to knock down OXTR binding in the NAc infected with the sg-RNA targeting OXTR, but not on the contralateral side infected with the sg-RNA targeting Lac Z (Figure 3G2). The LacZ hemisphere responded electrophysiologically to TGOT application with a potentiation of the eEPSP_f_ amplitude, as expected. The response of the OXTR knocked-down hemisphere was significantly reduced, producing a significant interaction (Figure 3H-I). Altogether, these data show that the activation of OXTR located on NAc cells is necessary for the OXT-induced potentiation of glutamatergic transmission into NAc, yet this potentiation likely arises from increased presynaptic release. This surprising result is in contrast to a previously described presynaptic mechanism for oxytocinergic modulation in mice (Dölen et al. 2013). Thus, we hypothesized that in pair bonded voles specifically, a retrograde signal may serve to link postsynaptic OXTR within the NAc to presynaptic modulation of excitatory transmission.

### CB1 receptor activation is necessary for OXT-dependent potentiation in pair bonded females

Endocannabinoids (eCB) are retrograde signaling molecules modulating the synaptic strength between cells (Castillo et al. 2012) and have been implicated in the electrophysiological (Ninan 2011; Hirasawa et al. 2004; Xiao, Priest, and Kozorovitskiy 2018) and behavioral (Wei et al. 2015) effects of OXT. Indeed, OXTR can be coupled to the intracellular signaling G-protein, Gq (Busnelli et al. 2013), leading to phospholipase C activation, a mechanism which can promote eCB production (Lu and Mackie 2016). To evaluate whether the OXT-dependent potentiation in Paired females relies on the activation of the eCB system, we performed electrophysiological experiments in field configuration and measured the effect of TGOT bath application in presence or absence of a CB1 receptor antagonist (AM4113, 10^−6^M). Using separate slices from the same animals as a control, we showed that AM4113 application prevented the TGOT-induced potentiation in Paired females (Figure 4A). Interestingly, this application also prevented the correlation between the effect of TGOT on the PPR and on the eEPSP_f_ amplitude (Figure 4B-C). Hence, the activation of the eCB system is necessary for the TGOT-induced potentiation observed in Paired females.

**Figure 4:**
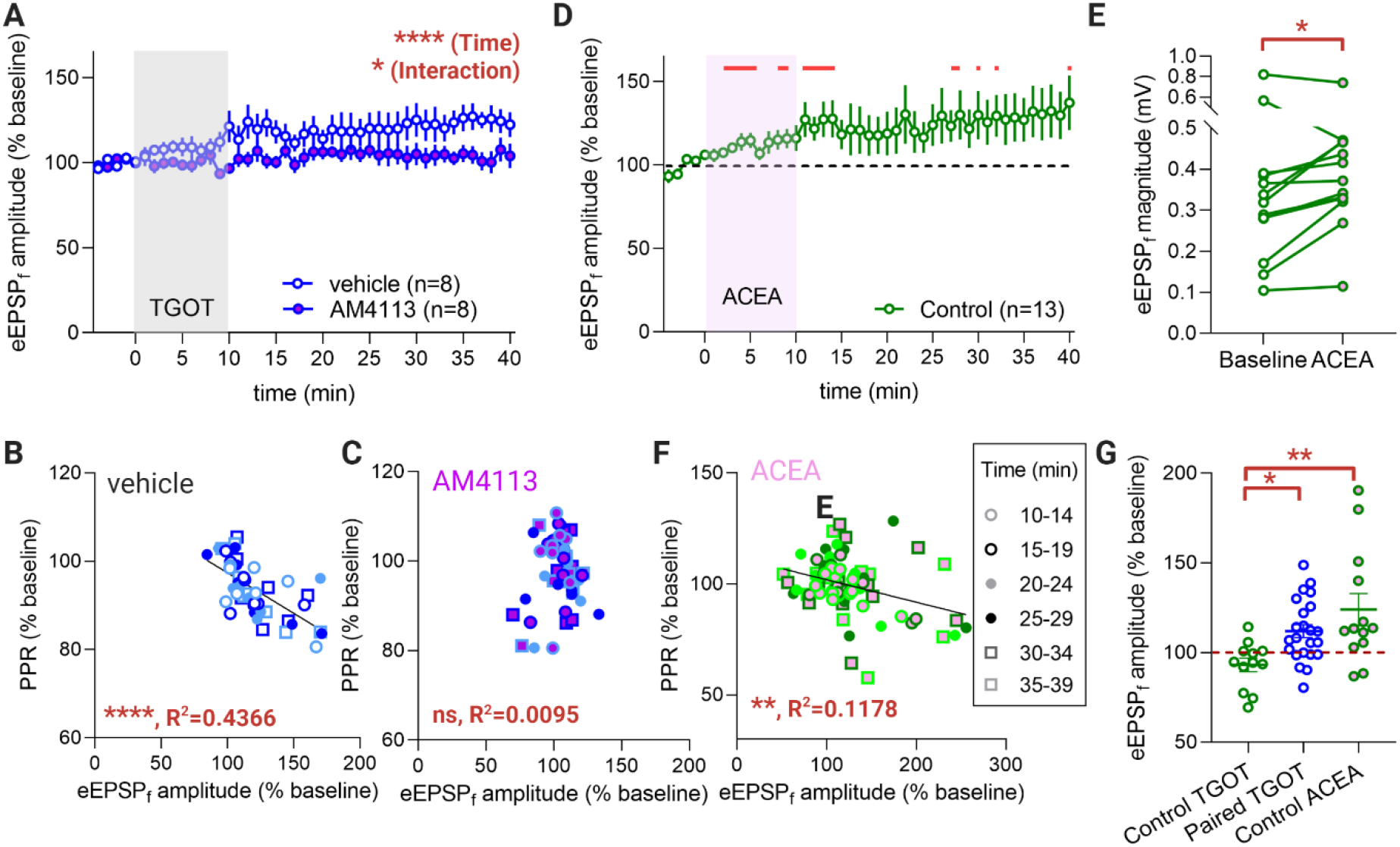
CB1 receptor activation is necessary for the TGOT-dependent potentiation. **(A)** Time course of the effect of 10min bath application of TGOT on evoked EPSP amplitude in pair bonded females in presence or absence of AM4113 (field recording). Data (mean ± SEM) expressed as % relative to baseline, n as indicated, one slice per condition for each animal and data Paired by experimental animal. Two-way ANOVA: effect of time F(44,308)=2.292, p<0.0001; effect of AM4113 F(1,7)=4.183, p=0.080; interaction F(44,304)=1.597, p=0.013. **(B)** Correlation between EPSP amplitude and Paired pulse ratio (PPR) in in field recordings from pair bonded animals after treatment with TGOT in absence of AM4113. Pearson correlation coefficient as indicated, p<0.0001. Linear regression: y=-0.1834x+115.7. **(C)** Correlation between EPSP amplitude and Paired pulse ratio (PPR) in in field recordings from pair bonded animals after treatment with TGOT in presence of AM4113. Pearson correlation coefficient as indicated, p=0.509. **(D)** Time course of the effect of 10min bath application of ACEA on evoked EPSP amplitude in Virgin females (field recording). Data (mean ± SEM) expressed as % relative to baseline, n as indicated, one slice per condition for each animal. One sample Wilcoxon test (compared to 100%), p<0.05 at min 2, 3, 4, 5, 8, 9, 11, 12, 13, 14, 27, 28, 30, 32 and 40. **(E)** Average over a 5min period (10 to 15min after the beginning of the application) of the effect of a 10min ACEA application on the evoked EPSP magnitude (absolute value of the amplitude) in NAc of Virgin animals. Wilcoxon test, p=0.0479. **(F)** Correlation between EPSP amplitude and Paired pulse ratio (PPR) in in field recordings from Virgin animals after treatment with ACEA. Pearson correlation coefficient as indicated, p=0.002. Linear regression: y=-0.1010x+112. **(G)** Average over a 5min period (10 to 15min after the beginning of the application) of the effect of 10min bath application of TGOT or ACEA on evoked EPSP amplitude (field recording). Data (mean ± SEM) expressed as % relative to baseline. One-way ANOVA: p=0.003, Holm-Sidak post-hoc tests: Virgin TGOT vs Paired TGOT: p=0.0363, Virgin TGOT vs Virgin ACEA: p=0.002, Paired TGOT vs Virgin ACEA: p=0.113.

### Pair bonding couples inherent eCB potentiation to OXTR signaling

Importantly, CB1 receptors are expressed in the NAc of sexually naive females (Simmons et al. 2021). The application of a CB1 receptor agonist, arachidonyl-2’-chloroethylamide (ACEA), produces a potentiation of the post-synaptic response to an electrical stimulation of NAc core in Virgin females (Figure 4D and E). The amplitude of the effect of ACEA on the eEPSP_f_ correlated with the effect of ACEA on the Paired pulse ratio (Figure 4F), confirming a presynaptic function. Interestingly, the amplitude of the response to ACEA in Virgin females is comparable to the amplitude of the response to TGOT in Paired females (Figure 4G). Thus, together our results suggest that in Virgin animals, OXTR activation is not coupled to a functioning CB1-mediated potentiation of excitatory transmission in the NAc, but pair bonding establishes this link between these neuromodulatory systems.

### Blocking CB1 receptor activation in the NAc of Paired females selectively increases partner rejection

The eCB-mediated electrophysiological effect of TGOT emerges only after socio-sexual experience. Although OXT presumably has many different effects, we wanted to determine whether disrupting the ability for eCB to induce this potentiation in NAc has any behavioral consequence for social interactions. We therefore implanted bilateral cannulas in the NAc core of ovariectomized females to locally deliver AM4113 (Figure 5A). We cohabitated estradiol-primed subjects for 24h with a male and then conducted a first SPT the next day. One day later, we administered subjects with a CB1 receptor blocker (or vehicle) in the NAc and conducted a second SPT (SPT2). During SPT2, irrespective of whether animals were injected with vehicle or AM4113, subjects spent more time on average near the partner than near the stranger (Figure 5B).

**Figure 5:**
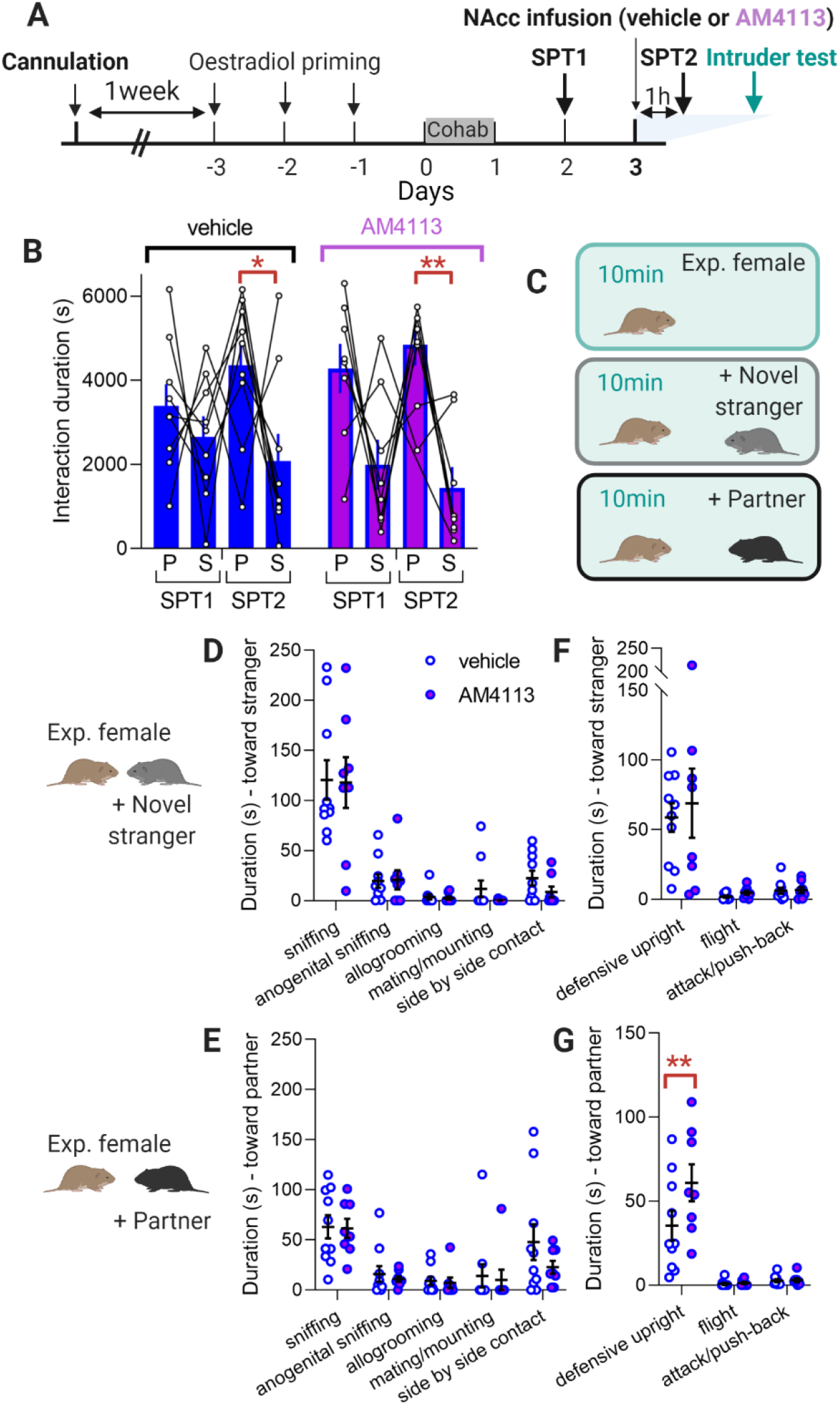
Blocking CB1 receptor activation in the nucleus accumbens selectively increases partner rejection but not stranger rejection. **(A)** Experimental timeline. **(B)** Duration of the interactions with partner (P) and stranger (S) during SPT1 and SPT2 in Paired females administered with vehicle or AM4113 in the NAc. Data (mean ± SEM), n=9 voles for SFP1 vehicle, 9 for SPF2 vehicle, 8 for SPF1 AM4113 and 8 for SPF2 AM4113. Three-way ANOVA: effect of treatment F(1,16)=0.004774, p=0.946; SPT1 vs SPT2 F(1,16)=0.082, p=0.778; partner vs stranger F(1,16)=20.84, p=0.0003; interaction treatment × SPT1 vs SPT2 F(1,16)=0.08146, p=0.779; interaction treatment × partner vs stranger F(1,16)=1.932, p=0.184; interaction SPT1 vs SPT2 × partner vs stranger F(1,12)=6.724, p=0.023 ; interaction treatment × SPT1 vs SPT2 × partner vs stranger F(1,12)=0.237, p=0.6352. Post-hoc Sidak test (partner vs stranger): SPT1 vehicle p=0.994, SPT2 vehicle p=0.047, SPT1 AM4113 p=0.091, SPT2 Paired p=0.003. **(C)** Experimental protocol – intruder test. **(D)** Duration of the affiliative behaviors displayed toward the novel male in voles administered with vehicle of AM4113 in the NAc. Data (mean ± SEM), n=10 vehicle-treated voles and 8 AM4113-treated voles. Two-way ANOVA: behavior F(4,64)=39.48, p<0.0001; treatment F(1,16)=0.4442, p=0.515; interaction F(4, 64)=0. 1804, p=0.948; subject F(16,64)=1. 468, p=0.140. Post hoc Sidak test (vehicle vs AM4113): sniffing p=0>9999; anogenital sniffing p>0.9999, allogrooming p>0.9999, mating/mounting p=0.962, side-by-side contact p=0.923. **(E)** Duration of the affiliative behaviors displayed toward the partner in voles administered with vehicle of AM4113 in the NAc. Data (mean ± SEM), n=10 vehicle-treated voles and 8 AM4113-treated voles. Two-way ANOVA: F(4,64)=11.15, p<0.0001; treatment F(1,16)=0.8955, p=0.358; interaction F(4,64)=0.5279, p=0.716; subject F(16,64)=1.669, p=0.076. Post hoc Sidak test (vehicle vs AM4113): sniffing p=0>9999; anogenital sniffing p=0.999, allogrooming p>0.9999, mating/mounting p=0.999, side-by-side contact p=0.373. **(F)** Duration of the aggressive/defensive behaviors displayed toward the novel male in voles administered with vehicle of AM4113 in the NAc. Data (mean ± SEM), n=10 vehicle-treated voles and 8 AM4113-treated voles. Two-way ANOVA: behavior F(2,32)=22.87, p<0.0001; treatment F(1,16)=0.2693, p=0.611; interaction F(2,32)=0.8797, p=0.880; subject F(16,32)=1.118, p=0.380. Post hoc Sidak test (vehicle vs AM4113): defensive upright p=0.862; flight p=0.997; attack/push-back p>0.9999. **(G)** Duration of the aggressive/defensive behaviors displayed toward the partner in voles administered with vehicle of AM4113 in the NAc. Data (mean ± SEM), n=10 vehicle-treated voles and 8 AM4113-treated voles. Two-way ANOVA: behavior F(2,32)=45.26, p<0.0001; treatment F(1,16)=3.161, p=0.094; interaction F(2,32)=3.345, p=0.048. Post hoc Sidak test (vehicle vs AM4113): defensive upright p=0.009; flight p>0.9999; attack/push-back p>0.9999.

Since the electrophysiological experiments above were performed after the pair bond was established, the eCB-dependent potentiation may naturally occur after pair bond formation and influence pair bond maintenance even in the absence of an effect on pair bond formation (Simmons, Singh, and Bales 2020). Furthermore, the NAc plays a role in action selection, so mechanisms acting at this time point may affect the nature of social interactions. To assess this, we performed intruder tests after SPT2. The female was first placed alone in her home-cage for 10 minutes, before a novel stranger male was introduced for 10 minute and then replaced by the partner for the final 10 minutes (Figure 5C). Since side by side contact was not influenced by CB1 receptor antagonist in a previous study (Simmons, Singh, and Bales 2020), we hypothesized that a CB1-dependent change in the quality of the social interactions would likely occur through a change in agonistic interactions rather than a change in pro-social interactions.

In the intruder test, affiliative and aggressive/defensive behaviors were scored blind to treatment group to evaluate the behavioral response of the female to the presence of a stranger male or the partner. No effect of the treatment was observed in affiliative behaviors performed by the female (Figure 5D-E) or the stimulus male (Supplemental Figure 3F-G)_-_, whether the female interacted with a stranger male or with her partner. However, for aggressive/defensive behaviors, there was an effect of treatment in the presence of the partner (Figure 5G). This was not the case in presence of the stranger (Figure 5F). Specifically, animals treated with AM4113 spent about twice as much time in a defensive upright posture when the partner was present, as compared to vehicle treated animals (Figure 5G), despite showing no difference in overall rearing behavior within the cage (Supplemental Figure 3A-B-C)

Hence, CB1 receptor activation occurs during a female’s social interaction with her partner to decrease the expression of a rejection-like, upright rearing response when the partner is present. Thus, this eCB-dependent mechanism would play an important role in defining the behavioral trajectory of social interactions by reinforcing a pair bond through a reduction of partner rejection.

## Discussion

Our study demonstrates that adult social experience, modeled here by pair bonding in female prairie voles, can alter the electrophysiological action of neuropeptides on a brain area, the NAc, important for social information processing and social salience. Activating NAc OXTRs induces divergent electrophysiological responses in female voles that are pair bonded versus same-sex littermate-housed, even when the OXTR agonist is delivered in exactly the same way (e.g., concentration, timing, application mode). For littermate-housed animals, it decreases the amplitude of the sEPSC (Figure 1C), but for animals in pair bonds, it increases the amplitude of the evoked activity (Figures 1E,F and 2E,F), suggesting distinct strategies for OXT to modify the signal-to-noise ratio in excitatory neurotransmission. In Paired prairie voles, OXTR agonists act on receptors located on NAc cells (Figures 3G,H) and lead to an eCB-mediated synaptic potentiation (Figure 4A) involving a presynaptic mechanism (Figure 4B,C). Blocking CB1 receptors after the pair bond is established specifically increases a partner-directed rejection behavior without affecting stranger-directed rejection (Figure 5D and 5E). Hence, the socio-sexual experience-contingent eCB-dependent oxytocinergic potentiation of NAc MSNs described here could influence the trajectory of social behavior and promote the maintenance of a pair bond.

To our knowledge, this is the first study showing that acute oxytocinergic modulation of excitatory transmission can be different depending on an adult’s social history. Building social relationships presumably involves neural changes in the steps leading from perception of another to the behavioral action towards that individual. Altered connectivity within neural networks is usually attributed to synaptic plasticity (Rutherford et al. 2020; Marlin and Froemke 2017) or state-dependent neuromodulation (e.g., release or receptor expression) (Marlin et al. 2015; Baxter et al. 2020), rather than changes in how neurotransmitters themselves work within the network. In fact, most studies of how OXT facilitates social behavior assume its mechanistic stability over state and time in adults (Marlin et al. 2015; Wei et al. 2015; Bartz et al. 2011). Variability in the effects of OXT across individuals are then often presumed to be due to individual differences in how much OXT is released (Alaerts et al. 2019; Parker et al. 2017), how many OXTR there are (King et al. 2016; Ross et al. 2009) or methylation status of the OXTR gene (Puglia et al. 2015; Puglia, Connelly, and Morris 2018; Andari et al. 2020). Here however, we showed that the way OXT engages cellular pathways to affect glutamatergic neurotransmission is context-dependent, even when the average strength of postsynaptic responses is the same (see Figure 1D and Supplemental Figure 1G and H).

We found that the electrophysiological effect of OXT application differs in Virgin and Paired voles, with a potentiation of excitation mediated through a coupling between OXTR and CB1 receptor activation emerging only in the latter group (Figure 4). The potentiation we observed was unexpected, since OXT in the NAc core (Dölen et al. 2013), ventral tegmental area (Xiao, Priest, and Kozorovitskiy 2018) or prefrontal cortex (Ninan 2011) typically induces long-term depression (LTD) of glutamatergic transmission. CB1 receptors, which are found presynaptically (Castillo et al. 2012) and expressed in the vole NAc (Simmons et al. 2021), are often implicated in such OXT-dependent depression of excitatory transmission (Ninan 2011; Hirasawa et al. 2004; Xiao, Priest, and Kozorovitskiy 2018). On the other hand, OXT-dependent potentiation, while observable when combined with an electrical induction paradigm (Ninan 2011; Yaseen et al. 2019; Fang, Quan, and Kaba 2008), has not previously been linked to the eCB system. Interestingly, even though eCB itself is usually associated with synaptic depression, its effects can be bi-directional depending on the concentration of eCB, the type of presynaptic neuron targeted (GABAergic vs glutamatergic), the involvement of astrocytes (Piette et al. 2020), and even the precise timing of presynaptic-to-postsynaptic firing (Cui, Perez, and Venance 2018). Thus, our finding of a pharmacologically induced, eCB-mediated, OXT-dependent potentiation in pair bonded females illustrates a new way for these neuromodulators to work together in the NAc.

The physiological effect of an OXT-eCB-based potentiation could allow upstream inputs to gain enhanced salience by amplifying excitatory transmission. The eCB system is generally engaged in naturally rewarding contexts (Parsons and Hurd 2015), such as food ingestion (Méndez-Díaz et al. 2012), physical exercise (Muguruza et al. 2019), and sexual activity (Canseco-Alba and Rodríguez-Manzo 2019). Given OXT’s well-documented involvement in social-specific behaviors (Shamay-Tsoory and Abu-Akel 2016; Marlin and Froemke 2017), the coupling in pair bonded female voles between OXTR and the eCB system may therefore reinforce the value of social interactions, which can be behaviorally mediated through such an association in mice (Wei et al. 2015). The potentiation of glutamatergic signaling carrying partner-specific information into the NAc core might then act to promote certain positively valanced social interactions.

Our conclusions fit within a working model (Figure 6) for how OXT, eCB and glutamatergic inputs into NAc may interact *in vivo* to adapt social responses to the context based on one’s social experience with the other individual. In virgins, social interactions that facilitate future pair bonding (e.g. mating), would release OXT, resulting in decreased amplitude of sEPSCs in NAc MSN, which may serve to decrease noise, facilitating linking the olfactory cues of the future partner with the reward system (Walum and Young 2018) (Figure 6 left). In females that are already pair bonded though, a potentiation of the glutamatergic transmission would arise from a coincidence of events: the activation of the OXT-eCB pathway along with the activity of upstream neurons reflecting the familiar partner (Figure 6 right).

**Figure 6:**
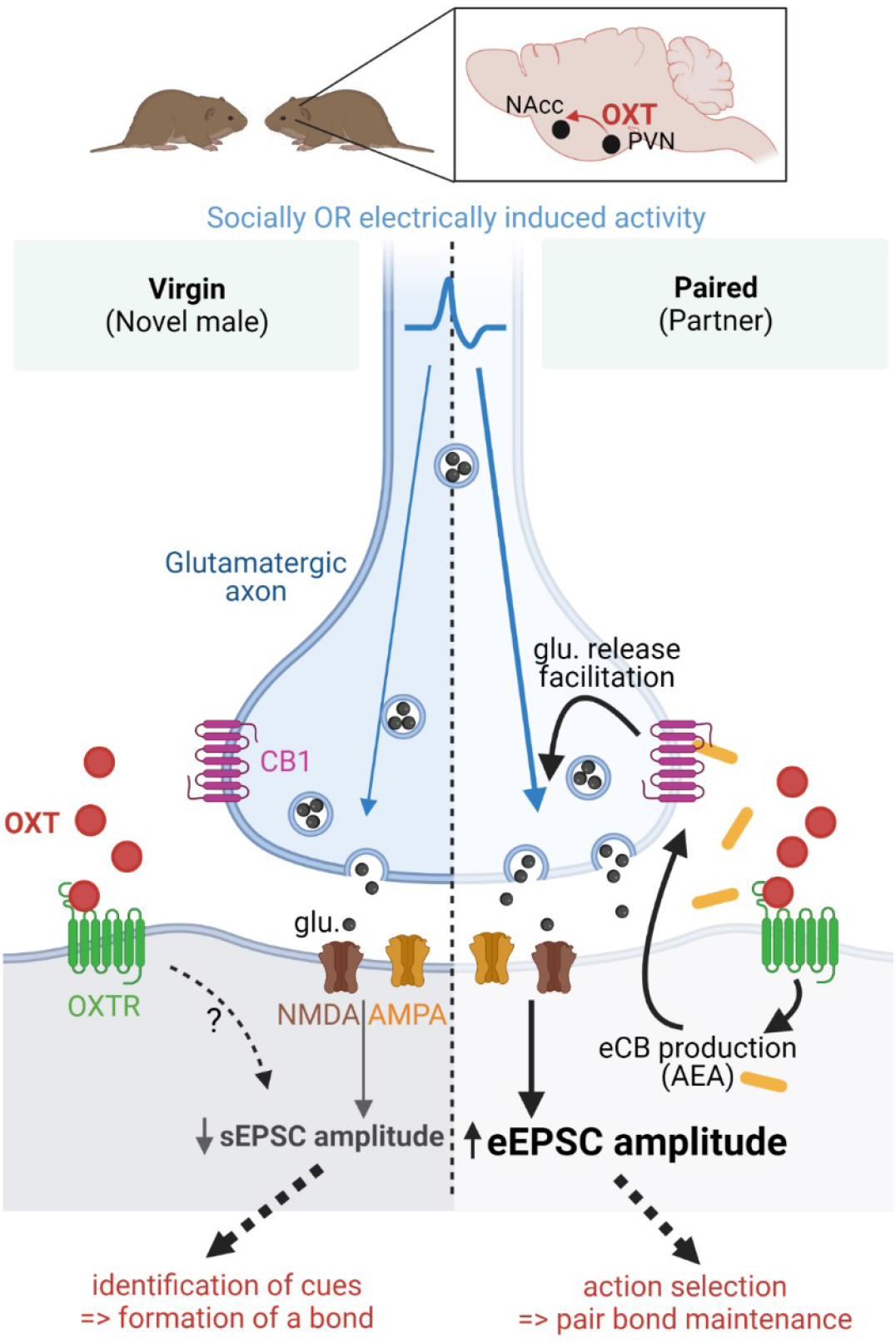
Working model. Model for the mechanism of OXT action on synaptic transmission in the nucleus accumbens of female prairie voles. AEA = anandamide, PVN = paraventricular nucleus, glu.= glutamate

Our behavioral results in pair bonded females suggest that only specific social interactions reveal the effects of NAc eCB. In particular, a subject’s defensive upright response to its partner – a form of social rejection – was increased by blocking NAc CB1 receptors (Figures 5E and F), without other effects on the subject’s general social preference (Figure 5B) or other social responses (Supplemental Figure 3). This finding is consistent with a recent pharmacological study in voles showing intraperitoneal AM4113 administration during cohabitation also did not affect partner preference as measured by side-by-side contact (Simmons, Singh, and Bales 2020), though defensive rejection was not assayed. In fact, subtle social interactions are not generally analyzed in pair bonding, and our results suggest methods to do so more broadly (Cande et al. 2018; Pereira, Shaevitz, and Murthy 2020) could be beneficial for understanding the specific role played by neuromodulatory systems and how their functions are coordinated to allow for the emergence of new behavioral repertoires.

To explain the context-dependent effect of the CB1 antagonist in pair bonded females in response to a novel and partner male, two non-exclusive mechanisms may play a role: OXT release could be higher in response to the familiar male as previously suggested (Ross et al. 2009; Lukas et al. 2013), or interacting with a familiar versus a novel conspecific could differentially activate regions upstream of the NAc (Okuyama et al. 2016). Importantly though, in our slice electrophysiology experiments, we controlled for both OXT concentration and presynaptic stimulation intensity to investigate whether the NAc also changes *how* it responds to this coincidence. We presume that for interactions with the partner but not novel male, OXTR activation leads to the local production of anandamide, a potent agonist of CB1 receptors. This facilitates the release of more glutamate upon activation of the pre-synaptic neurons (Figure 1E,F and 2E,F), while leaving spontaneous synaptic transmission unchanged (Figure 1B and C). Such an OXT-dependent mechanism could help filter salient inputs to the NAc – similar to how OXT enhances the signal-to-noise ratio in other brain areas (Owen et al. 2013; Oettl et al. 2016; Froemke and Young 2021) by a mechanism that here involves the modulation of another neuromodulatory system: the eCB system. Furthermore, we hereby demonstrate that such an OXT-dependent modulation of glutamatergic transmission can depend on the context defined by the socio-sexual experience.

*In vivo*, the putative mechanism described above would potentiate the effect of socially induced electrical presynaptic activity, leading to the inhibition of rejection behavior when the partner is present. In Virgins, this inhibition would not happen and interactions would be more agonistic (Lee et al. 2019; Bowler, Cushing, and Sue Carter 2002). This working model may help to conceptualize the mechanisms underlying context-dependent behavioral differences resulting from exogenous OXT administration in humans (Bartz et al. 2011). Indeed, the effect of intranasal OXT depends on the partner status of the subject: it keeps men in a monogamous relationship from being too close to a new attractive woman without affecting single men (Scheele et al. 2012). OXT’s effects further depend on the interrelationship between a subject and the social stimuli: it increases the perceived attractiveness of one’s partner but not of other women, even if they are familiar (Scheele et al. 2013).

Interestingly, the authors of that study also reported that in men seeing their partner, OXT increases NAc’s response to the partner pictures compared to familiar and unfamiliar women.

Finally, it has long been believed that OXT acts in the NAc core to first establish the bond (Johnson et al. 2017; Walum and Young 2018), and then to increase the relative value of the partner, thus promoting the maintenance of the bond. The results presented here suggest that in both aspects of pair bonding, OXT could act through a modulation of the signal-over-noise ratio (i.e. decreasing the noise during the first social interaction and increasing the signal after the bond is established). Importantly, socio-sexual experience appears to condition the reinforcing properties of the OXT system through its association with the eCB system. This could sustain behavioral changes that are happening as the social relationship with a given individual unfolds, and alterations in the described mechanism could negatively affect the trajectory of social relationships. By showing that context influences OXT’s mode of action in the NAc, the present study offers a potential new explanation for inconsistent outcomes of OXT’s effects in human studies (Alvares, Quintana, and Whitehouse 2017). Indeed, while inter-individual differences were identified as the likely source of variability in clinical trial outcome, it is usually attributed to variation in the intrinsic properties of the OXT system (Andari, Hurlemann, and Young 2018; Kosaka et al. 2016). Here, we point out that a complementary explanation – social experience itself influencing the mode of action of OXT – should motivate the factoring in of social history when considering the effect of OXT treatments.

## Methods

### Animals

All experiments were performed in accordance to the guidelines and approved by the Emory University Institutional Animal Care and Use Committee. Prairie voles bred at Emory University originated from field caught specimens from Illinois, USA. Weaning was performed at postnatal day 20-23 and animals were house-grouped (2 to 3 voles per cage) in same-sex sibling cages.

Prairie voles between 2 and 6 months old were used for all the experiments. Animals were housed under a 14:10 hour light/dark cycle, temperature was maintained at 68-72 degrees Fahrenheit and humidity at 40-60%. Food (Lab Rabbit Diet HF #5326, LabDiet) and water were given ad libitum. Enrichment consisting of cotton pieces used to build a nest were added to cages. When animals were isolated, supplemental enrichment material (Nylabones, Bioserve) was added to the cage. All females used were ovariectomized prior to the beginning of the experiments.

### Slice electrophysiology

#### Slicing

After isoflurane anesthesia followed by quick decapitation, brains were extracted and 300μm thick coronal slices were sliced using a Leica VTS-1000 vibrating-blade microtome in ice-cold slicing solution (in mM: 130 NaCl, 3.5 KCl, 1.1 KH_2_PO_4_, 6 MgCl_2_, 1 CaCl_2_, 10 glucose, 2 kynurenate, 30 NaHCO_3_, bubbled with 95% O_2_ and 5%CO_2_). Slices were transferred to a 32°C slicing solution for 1h and then to room-temperature artificial cerebro-spinal fluid (ACSF) (in mM: 130 NaCl, 3.5 KCl, 1.1 KH_2_PO_4_, 1.3 MgCl_2_, 2.5 CaCl_2_, 10 glucose, 0.4 ascorbate, 0.8 thiourea, 2 Na pyruvate, 30 NaHCO_3_, 95% O_2_ and 5%CO_2_).

#### Recordings

Slices were transferred to the recording chamber mounted on the fixed stage of a Leica DM6000 FS microscope (Leica Miscrosystems Inc. Bannackburn, IL) and perfused at ≈2mL/min with ACSF maintained at 32°C. Recordings were performed using MultiClamp700B amplifier in conjunction with pClamp 10.2 software and a DigiData 1322A (Axon Instruments).

Borosilicate glass patch electrodes (WPI, Sarasota, FL, USA) were used to acquire whole-cell patch-clamp or field recordings in the NAc core. For whole cell recordings, electrodes had a resistance of 4–7 MΩ, were filled with a “patch solution” (in mM: 130 K-gluconate, 2 KCl, 10 HEPES, 3 MgCl2, 2 K-ATP, 0.2 Na-GTP, 5 phosphocreatine, adjusted to pH 7.3 with KOH). Pipette offset and capacitance were automatically compensated for using MultiClamp command software (Molecular Devices). For field recordings, electrodes were filled with ACSF.

A concentric bipolar stimulation electrode was placed near the recording electrode and moved inside the NAc core until a signal was perceived by the recording electrode. The intensity of the stimulation was defined at the beginning of each recording such as it induces 50% of the maximum evoked response. A pair of stimulations with 50ms interval were applied every 30s during the recording. The values for eEPSC and eEPSP_f_ were obtained in response to the first stimulation. The influence of TGOT on spontaneous EPSCs was measured between 1 and 5s after the stimulation.

Recordings from fast spiking interneurons were excluded based on their electrophysiological properties so that the results presented here were obtained exclusively from MSNs.

#### Pharmacology

Pharmacological compounds were diluted in ACSF and bath applied. The concentration used are the following: TGOT 10^−7^M (Sigma-Aldrich), AM4113 10^−6^M (Sigma-Aldrich), DNQX 2×10^−5^M (Sigma-Aldrich), ACEA 10^−7^M (Tocris). TGOT was applied for 10min. AM4113 was applied for at least 10min before the application of TGOT and was maintained for the duration of the recording. DNQX application lasted for 10min and was performed at the end of the recording.

### Behavior: Social preference test (SPT)

All females were ovariectomized and primed for 3 days with subcutaneous administration of estradiol benzoate (17-β-Estradiol-3-Benzoate, Fisher Scientific, 2 μg dissolved in sesame oil starting 3 days prior to experiments). The next day, a SPT could be performed before animals were cohabitated with a same sex littermate or with one of the males used in the SPT for 24h. Previous studies showed that 24 h cohabitation with a male induces a strong pair bond as indicated by more time spent huddling with the partner compared to the stranger in a partner preference test (Williams, Catania, and Carter 1992). Animals were then isolated until euthanasia and brain extraction which occurred 2 days later except for a potential second SPT test which would be performed on the day prior to euthanasia.

For the SPT, two males of similar age and weight were used as stimuli. One of them was randomly assigned as the future partner and was cohabitated with the experimental female after the test. The experimental arena consisted of a rectangular arena (30×8 inches). Two metallic cups (4 inches diameter and 4 inches height) constituted of metal bars were placed on each sides of the arena. Males were placed under these cups shortly before the female was introduced in the middle of the arena. The female was given 2h to freely explore the arena and/or the cups. When 2 SPT were performed, the same males were used as stimuli. For experiments comparing Virgin and Paired voles, 2 littermate females were used for each experiment and were tested with the same 2 males.

IdTracker was used to track the center of mass of the experimental subjects (Pérez-Escudero et al. 2014). A custom R code was then used to calculate the duration spent in a circle of a radius equal to 3 times the radius of the cup for each cup thus defining the time spent in the “partner zone” and the time spent in the “stranger zone”. Preference for the partner was calculated as follow: time in the partner zone (in s) / time in the stranger zone (in s).

### Surgeries

Anesthesia induction was achieved with 3% Isoflurane and maintained with 1-3% Isoflurane during the surgery. Meloxicam 2mg/kg was subcutaneously administered before the surgery begins and for 2 days following the surgery. Animals were monitored daily and were given at least 1 week to recover after surgery before experiments began. Anterior/posterior coordinates were referenced to Bregma, and dorsal/ventral coordinates were referenced to the top of the skull. Animals were placed in a stereotaxic frame and ear bars coated with Lidocaine were used to stabilize the head.

#### • Virus injection

500nL of viruses were injected in the NAc core (10° angle, +1.9mm AP, +/−2.6mm ML, −4.5mm DV) using a 10μL syringe (nanofil, World Precision Instrument, USA) with NanoFil Needle (NF33BV-2, World Precision Instruments, USA) at a rate of 100nL/min (Micro4 pump, World Precision Instruments, USA).

#### • Cannula implantations

Bilateral guide cannulas (26G – P1 Technologies, USA) heading to the NAc core were implanted (AP +1.5mm, ML +/−1.5cm, DV −4.3cm). Dental cement (Glass Ionomere cement – Harvard Apparatus, USA) and sutures (Ethicon J493G – ShopMedVet, USA) were used to secure the implant.

### Intra-cerebral infusions

AM4113 was diluted in ACSF and used at a final concentration of 20μM with 0.1% DMSO. The vehicle was ACSF with 0.1% DMSO.

Internal cannulas (P1 Technologies, USA) with a projection of 0.2mm were used for the injections. They were connected through 2 different tubing to two 1 μL Hamilton syringes (P1 Technologies, USA) controlled by a microinjector pumps (micro 4, World Precision Instrument, USA). Injections of 500nL/hemisphere were performed at 100 nL/min in isoflurane anesthetized voles. Internal cannulas were left in place 5min after the end of the injection to allow for diffusion. Experiments were started 1h after the beginning of the injection. Positions of cannulas were verified *post mortem*.

### AAV-mediated Crispr Knock-down of OXTR

AAV9-particles were synthesized usingpAAV-U6-sgRNA-CMV-eGFP and pAAV-RSV-spCas9 (Addgene plasmids #85451 and 85450, kindly gifted by Hetian Lei) , pAAV9-SPAKFA (Penn Vector Core, PA, USA) and pAAV/Ad (ATCC, VA, USA), ). sgRNA sequences cloned into pAAV-U6-sgRNA-CMV –eGFP were the following: sgRNA(OXTR): 5’-GCTGCGGTGGCCCGGCTGTG-3’, sgRNA(LacZ): 5’-GTGAGCGAGTAACAACCCGT-3’. AAV9-particles were produced in HEK293T cells and purified with AVB-affinity chromatography (Wang et al. 2015). 3 virus were generated: AAV9-U6-sgRNA(Oxtr)-CMV-eGFP (3.0×10^10 genomic copies/μl), AAV9-U6-sgRNA(LacZ)-CMV-eGFP (3.0×10^10 genomic copies/μl) and AAV9-RSV-spCas9 (1.5×10^10 genomic copies/μl) and used as previously described (Boender and Young 2020). A minimum of 4 weeks was allowed for virus expression before the experiments started.

### OXTR autoradiography

Autoradiography was performed as previously described (Ross et al. 2009).

### Statistics

Data are shown as means±standard error of the mean (SEM). In testing statistical significance, a was set at 0.05. The n for each data set is indicated in figures and/or legends. All data were averaged per experimental groups. Data were considered non-parametric when sample size was moderate. We used factorial Analysis of Variance (ANOVA) to compare multiple groups followed by post hoc comparisons with Sidak tests. All statistical analysis was performed with GraphPad Prism 9.

## Authors contributions

AMB, LJY and RCL conceptualized the project. AMB, SA and PL carried out the experiments. SA, PL and GJB designed the methodology to analyze the location of the animals. AJB designed and produced the CRISPR viruses. JG and HZ provided feedback for the electrophysiology experiments. AMB and RCL wrote the first draft and all the authors edited, reviewed and approved the final manuscript.

## Acknowledgements

This work was funded by NIH grant R01MH115831 (GJB and RCL) and Conte Center grant P50MH100023 (LJY and RCL). We thank Lorra Julian and the Yerkes animal care and veterinary staff for vole husbandry and care, and Dr. Jamie LaPrairie for her assistance. Figures and schematics were created using Biorender.

