## Supplemental figures for "Social experience alters oxytocinergic modulation in the nucleus accumbens of female prairie voles"

**Supplemental Figure 1 (refers to Figure 1): Basal properties of NAc core neurons do not depend on the pair bonding status of the experimental animal**

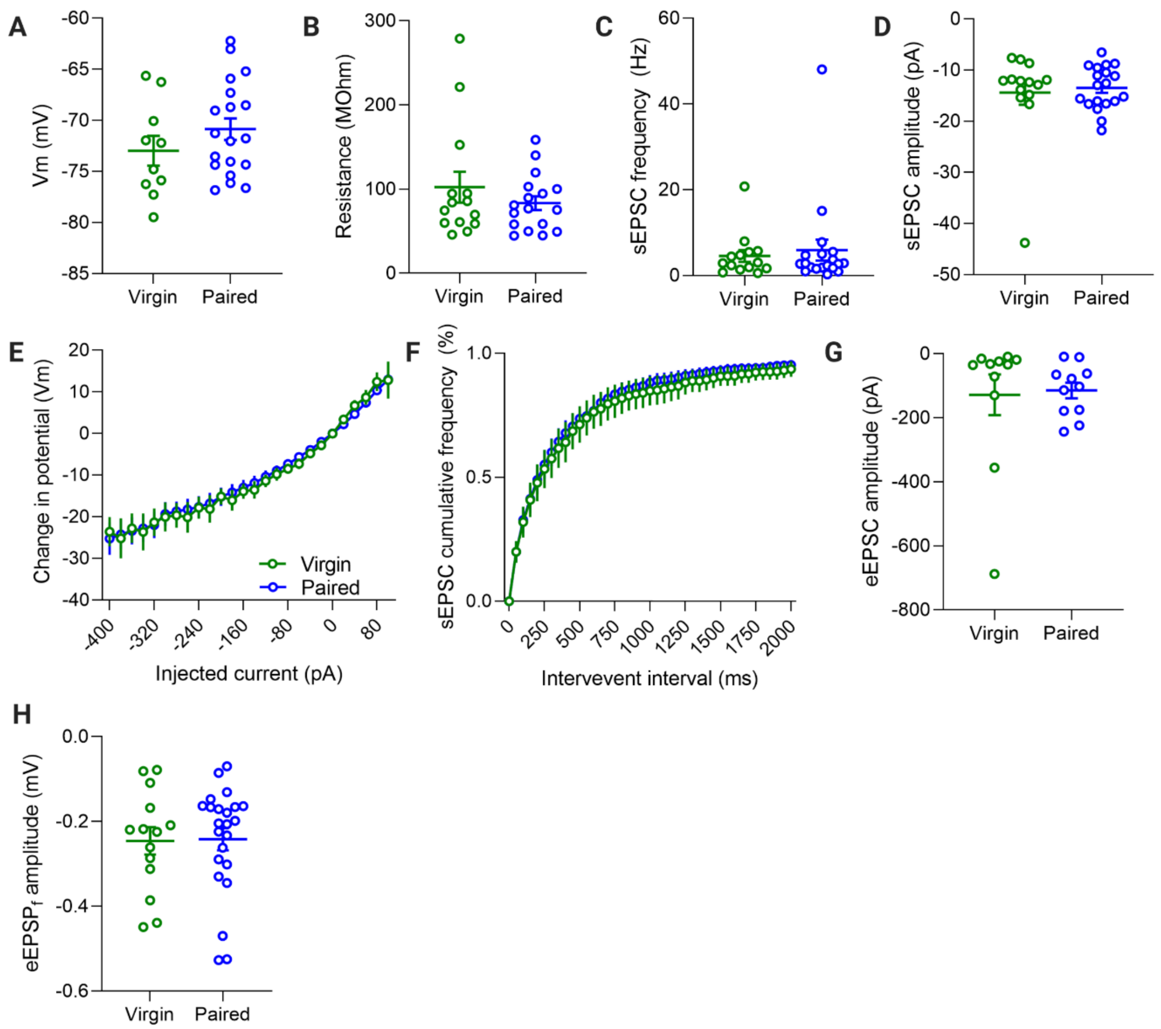


**(A)** Resting membrane potential of prairie vole neurons. Data (mean ± SEM) from n=10 neurons from Virgin voles and 19 neurons from Paired voles. Mann-Whitney test p=0.228. **(B)** Membrane resistance of prairie vole neurons. Data (mean ± SEM) from n=14 neurons from Virgin voles and 17 neurons from Paired voles. Mann-Whitney test p=0.681. **(C)** Frequency of the spontaneous EPSCs recorded from prairie vole neurons. Data (mean ± SEM) from n=14 neurons from Virgin voles and 19 neurons from Paired voles. Mann-Whitney test p=0.822. **(D)** Amplitude of the spontaneous EPSCs recorded from prairie vole neurons. Data (mean ± SEM) from n=14 neurons from Virgin voles and 19 neurons from Paired voles. Mann-Whitney test p=0.602. **(E)** Input-output curves of neurons recorded from Virgin and Paired voles. Data (mean ± SEM) from n=14 neurons from Virgin voles and 19 neurons from Paired voles. 2-way ANOVA: current intensity F(25,648)=80.29, p<0.0001 ; Social exp. F(1,31)=0.01611, p=0.900 ; interaction F(25,648)=0.1726, p>0.9999. **(F)** Cumulative distribution of the spontaneous EPSC as a function of their interevent interval in Virgin and pair bonded voles. Data (mean ± SEM) from n=14 neurons from Virgin voles and 19 neurons from Paired voles. 2-way ANOVA: interevent interval F(40,1240)=219.6, p<0.0001 ; Social exp. F(1,31)=0.1510, p=0.700 ; interaction F(40,1240)=0.1078, p>0.9999, cell F(31,1240)=154.8, p<0.0001. **(G)** Amplitude of evoked EPSC in baseline in whole cell recording before application of TGOT in Virgin and pair bonded voles. Data (mean ± SEM) from n=11 slices from Virgin voles and 11 slices from Paired voles. Mann Whitney test, p=0.365. **(H)** Amplitude of evoked EPSP in baseline in field recording before application of TGOT in Virgin and pair bonded voles. Data (mean ± SEM) from n=14 slices from Virgin voles and 23 slices from Paired voles. Mann Whitney test, p=0.699.

**Supplemental Figure 2 (refers to Figure 5): Location of the cannulas**

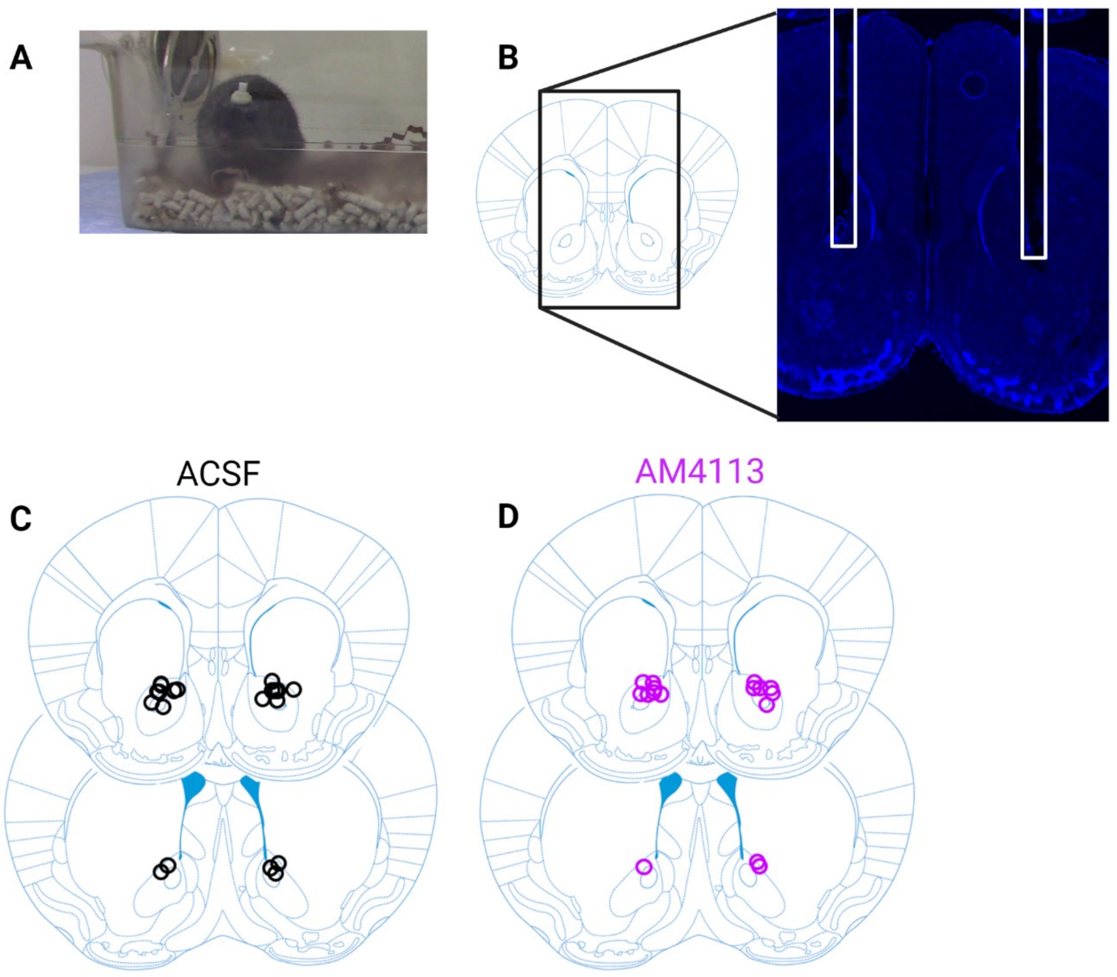


**(A)** Picture of a prairie vole implanted with the bilateral cannulas. **(B)** Representative picture of the images used to locate the cannulas *post-mortem*. Cannula location, indicated in white, is extrapolated from the lesion (blue = DAPI staining of the nuclei). **(C)** Location of the cannulas in ACSF-treated animals (n=10 animals). **(D)** Location of the cannulas in AM4113-treated animals (n=8 animals)

**Supplemental Figure 3 (refers to Figure 5): Effect of CB1 receptor antagonist administration on affiliative and neutral behaviors and on the behavior displayed following sniffing by the stranger**

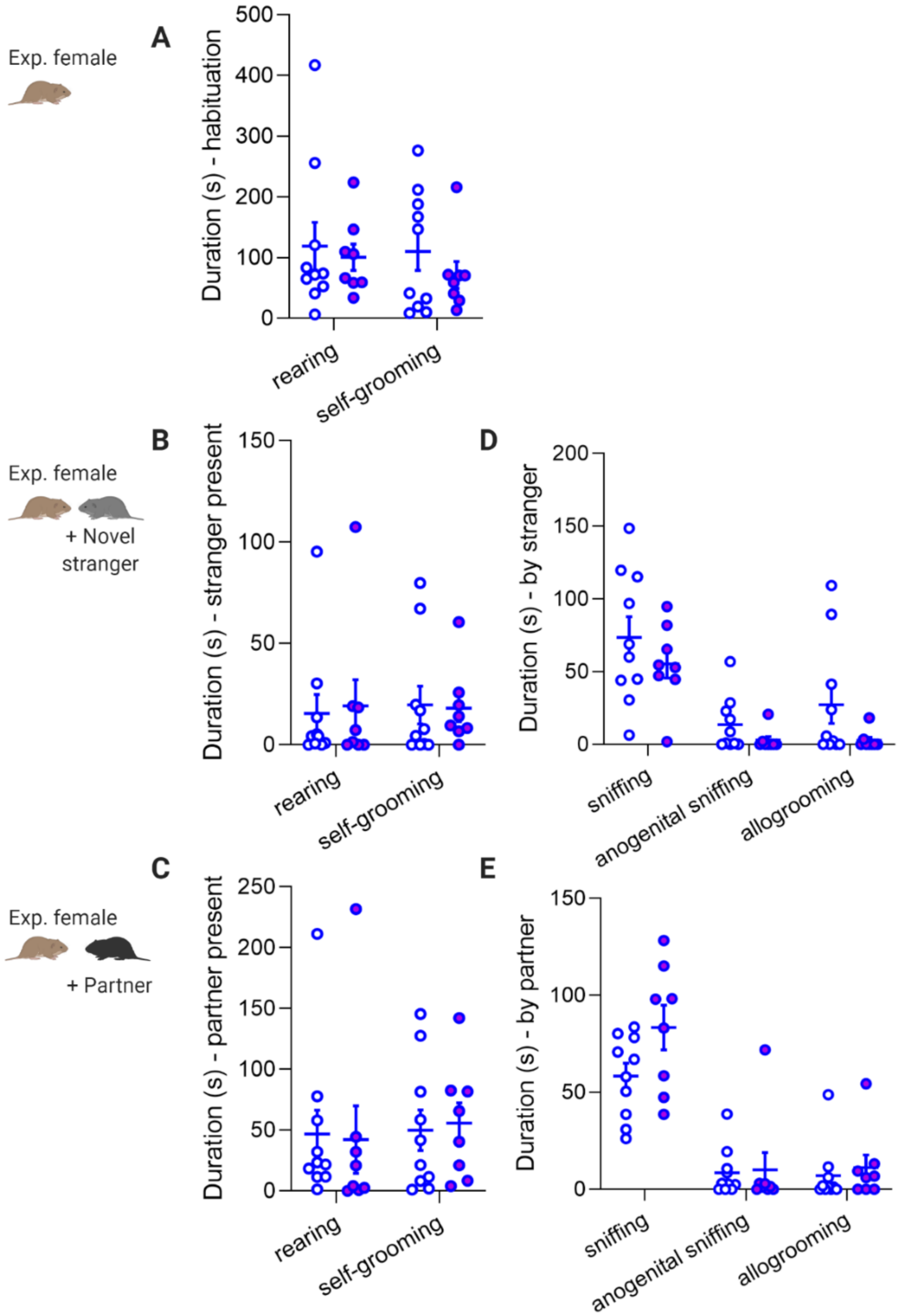


**(A)** Duration of the neutral behaviors exhibited by the experimental animal during the habituation period. Data (mean ± SEM) from n=10 vehicle-treated and n=8 AM4113-treated voles. Two-way ANOVA: behavior F(1,16)=0.2388, p=0.632; treatment F(1,16)=1.670, p=0.215; interaction F(1,16)=0.0704, p=0.794; subject F(16,16)=0.3294, p=0.984. Post hoc Sidak test (vehicle vs AM4113): rearing p=0.899; self-grooming p=0.627. **(B)** Duration of the neutral behaviors exhibited by the experimental animal when the stranger male is present. Data (mean ± SEM) from n=10 vehicle-treated and n=8 AM4113-treated voles. Two-way ANOVA: behavior F(1,16)=0.0211, p=0.886; treatment F(1,16)=0.01352, p=0.909; interaction F(1,16)=0.06413, p=0.803; subject F(16,16)=0.773, p=0.6934. Post hoc Sidak test (vehicle vs AM4113): rearing p=0.956; self-grooming p=0.992. **(C)** Duration of the neutral behaviors exhibited by the experimental animal when the partner male is present. Data (mean ± SEM) from n=10 vehicle-treated and n=8 AM4113-treated voles. Two-way ANOVA: behavior F(1,16)=0.1890, p=0.669; treatment F(1,16)=0.0008359, p=0.977; interaction F(1,16)=0.07354, p=0.790; subject F(16,16)=1.207, p=0.356. Post hoc Sidak test (vehicle vs AM4113): rearing p=0.984; self-grooming p=0.974. **(D)** Duration of the affiliative behaviors displayed by the stranger male. Data (mean ± SEM) from n=10 vehicle-treated and n=8 AM4113-treated voles. Two-way ANOVA: behavior $F(2,32)= 28.25$, $p<0.0001$; treatment $F(1,16)= 2.896$, $p=0.108$; interaction $F(2, 32)= 0.3535$, $p=0. 705$; subject $F(16,32)= 2.452$, $p=0.015$. Post hoc Sidak test (vehicle vs AM4113): sniffing $p=0.495$; anogenital sniffing $p=0.8332$; allogrooming $p=0.243$. **(E)** Duration of the affiliative behaviors displayed by the partner male. Data (mean ± SEM) from n=10 vehicle-treated and n=8 AM4113-treated voles. Two-way ANOVA: behavior $F(2,32)=59.12$, $p<0.0001$; treatment $F(1,16)=2.404$, $p=0.141$; interaction $F(2,32)=1.932$, $p=0.161$; subject $F(16,32)= 1.515$, $p=0.155$. Post hoc Sidak test (vehicle vs AM4113): sniffing $p=0.048$; anogenital sniffing $p=0.998$; allogrooming $p=0.967$.
